# Functional ultrasound imaging through a human cranial window for mesoscopic mapping of motor effector encoding within the sensorimotor cortex

**DOI:** 10.64898/2026.07.03.735688

**Authors:** Lydia J. Lin, Thierri Callier, Baptiste Heiles, Kelsie Pejsa, Charles Y. Liu, Mikhail G. Shapiro, Richard A. Andersen

**Author notes:** **Corresponding author:** Correspondence should be addressed to L.J.L.

## Abstract

Understanding movement encoding within human cortical circuits has been essential for advancing brain computer interfaces (BCIs). However, there are limited minimally invasive, high resolution neurorecording methods sensitive enough to detect single-trial movement-correlated neural activity. Functional ultrasound imaging (fUSI) provides submillimeter spatial resolution of deep cortical tissue with high sensitivity and, when paired with acoustically transparent skull implants, enables transcutaneous recording of human neurovascular changes. Prior studies have used fUSI in participants with acoustically transparent skull implants for on-off task mapping and decoding. Here, we demonstrate fUSI’s ability to reliably resolve multi-body-part and single digit movement encoding within the primary sensorimotor cortex in a participant with an acoustically transparent skull implant. We obtained fine-grained mappings of individual effector representation that were consistent with classic somatotopy for both multi-body-part and single digit movement. We were able to resolve single-trial event-related activity, enabling single-trial decoding of both conditions. Analysis of voxels important for decoding suggested differential encoding of single digit movement information across the different Brodmann areas. Finally, we show that these patterns can be approximated across different sessions, allowing for cross session decoding. These results establish that fUSI can reliably delineate somatotopically organized motor representations at submillimeter resolution, bridging a critical gap between invasive electrophysiology and noninvasive hemodynamic imaging in a human subject.

## Main

Actions such as drinking from a glass of water are considered simple. Yet, in reality, human motor function is composed of a combination of complex muscle movements coordinated from the top-down by various neural mechanisms. Neurological injury and disease such as spinal cord injury and amyotrophic lateral sclerosis are debilitating conditions that impair these critical functions, significantly impacting patients’ daily living. Understanding the neural mechanisms of how individuals generate movement through steady advances in novel neurotechnology have been essential in improving these patients’ quality of life.

Brain computer interfaces (BCIs) leverage neurorecording techniques to help such patients achieve greater independence by predicting intended behavior from neural activity to control assistive devices. However, most current BCIs rely on electrophysiology, which requires highly invasive brain surgery for subdural electrode implantation. These electrodes are restricted to a small, superficial region of the brain, typically 4×4 mm with up to 1.5 mm in depth for Utah arrays, and have limited sampling density^1^. Electrocorticography (ECoG) provides a larger field of view but is similarly invasively implanted subdurally and still only provides information from superficial brain regions^2,3^. Additionally, electrode-based methods are prone to degradation and recording quality deterioration over time^1,4–8^. In contrast, conventional non-invasive imaging techniques like fMRI and EEG offer a large field of view without requiring surgical intervention. Yet they lack sufficient spatial resolution for high performance BCI use^9^, with fMRI having a typical spatial resolution of 3-4 mm - ∼1 mm for 7T magnets – and EEG having a resolution of > 10-20 mm^10^. fMRI, specifically, is limited in the behavior that can be observed during imaging due to the large, enclosing machinery required for imaging.

Functional ultrasound imaging (fUSI) is an increasingly popular neuroimaging technique that balances these tradeoffs by imaging from outside the dura with high sensitivity and high spatial resolution while maintaining a large field of view. fUSI can detect transient changes in cerebral blood volume (CBV) at submillimeter spatial resolution for depths up to 4.9 cm, offering a mesoscopic view of neural activity across large subcortical brain regions that was previously not possible^11–13^. fUSI probes are small and lightweight, with our probe measuring ∼12 x 51 x 30 mm (WxLxH). Though fUSI probes are typically connected to desktop-sized acquisition systems, limiting movement to within the reach of the probe, this allows for high resolution recording of neural activity during behavior in more natural settings compared to other noninvasive neuroimaging modalities. This makes fUSI a well-suited tool to study human behavior. It has been used to image freely moving rodents^14–16^, NHPs during reach and saccade tasks^17–19^, and humans during guitar-playing^13^ and walking^20^. Additionally, fUSI planes are highly stable at the mesoscopic level, allowing for imaging of similar neuronal populations across sessions^19^. Because fUSI can image from outside of the dura, there is reduced risk of tissue damage and probe degradation, making fUSI an ideal candidate for minimally invasive chronic imaging (Fig. 1a). This has broad applications for long-term neural monitoring of human behavior and, beyond this, as a minimally invasive alternative for BCI, with prior studies already demonstrating fUSI-BCI capabilities in NHPs^17,18^.

**Fig. 1:**
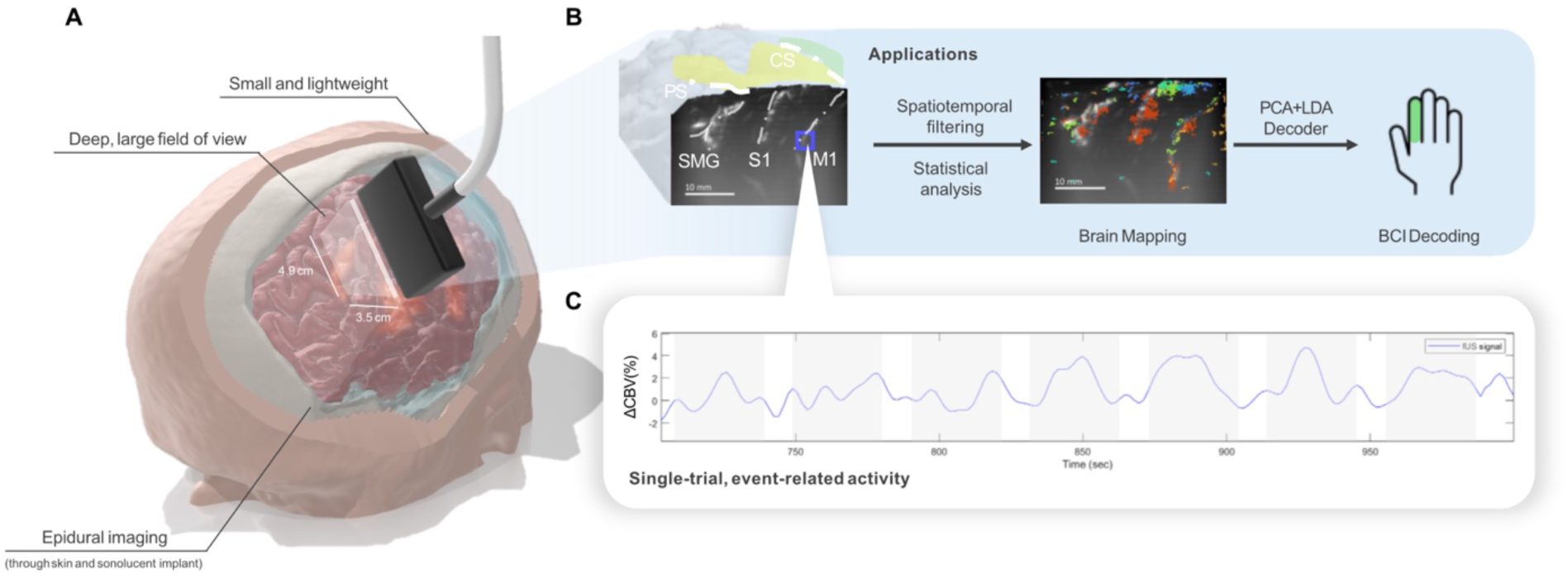
fUSI general overview and decoding pipeline. A) Simulated image of fUSI transducer recording brain activity (highlighted regions of brain) through an acoustically transparent cranial implant. fUSI is a small and portable neuroimaging modality that can image epidurally or from above the skin when paired with an acoustically transparent cranial implant, providing a deep, large field of view (3.5 cm W x 4.9 cm H) compared to electrophysiology. B) fUSI images can be preprocessed using spatiotemporal filtering to generate maps of task-correlated activity and decode behavior using neural signal. The displayed fUSI plane includes regions of M1, S1, and SMG and was used for all later analyses. C) Average fUSI signal over time, represented by percent change in cerebral blood volume (CBV), within the highlighted ROI. Grey, shaded boxes indicate movement execution periods.

However, fUSI currently is unable to image through the skull due to skull aberration and signal attenuation^21^. Because of this, fUSI requires a portion of the skull to be surgically removed through craniotomy and replaced with acoustically transparent material through cranioplasty^13^. For patients that require cranioplasty after neurosurgical operations, acoustically transparent implant materials such as polymethyl methacrylate (PMMA) and polyether ether ketone (PEEK) provide an opportunity to both clinically and scientifically better understand the brain without needing additional invasive procedures^22^. Acoustically transparent cranial implants allow for non-invasive transcranial ultrasonic imaging and have already been used for several medical and scientific applications. These include bedside ultrasonic monitoring of posttraumatic hydrocephalus after traumatic brain injury (TBI)^23^ and postoperative bypass patency as well as functional imaging in human participants during scientific behavioral tasks^13,20^. Thus, acoustically transparent cranial implants have enabled the expansion of fUSI to human clinical and scientific applications, paving the way for our current study.

Previously, it has been shown that fUSI can be used to accurately detect task state in a human participant through an acoustically transparent cranial implant^13^. In this study, we go further and demonstrate that fUSI can be used to map and decode multi-effector information from the sensorimotor cortex through an acoustically transparent cranial implant in the same participant, a proof of concept of fUSI’s potential as a neuroscientific tool and minimally invasive alternative for BCIs (Fig. 1). We first demonstrate the reliability of fUSI to accurately detect (Fig. 2) and distinguish differential movement activity over long periods of time by mapping out multi-body-part and then single digit movement activity within the sensorimotor cortex (Fig. 3,4). Using this information, we demonstrate fUSI’s ability to detect fine-grained differences in neural representation for successful decoding of motor effector information at significantly above chance levels (Fig. 5). We then examine the specific contributions of various Brodmann Areas (BA) to motor effector representation decoding (Fig. 6). Finally, we show that similar mappings can be identified across sessions recorded on different days, allowing for cross-session decoding (Fig. 7).

**Fig. 2:**
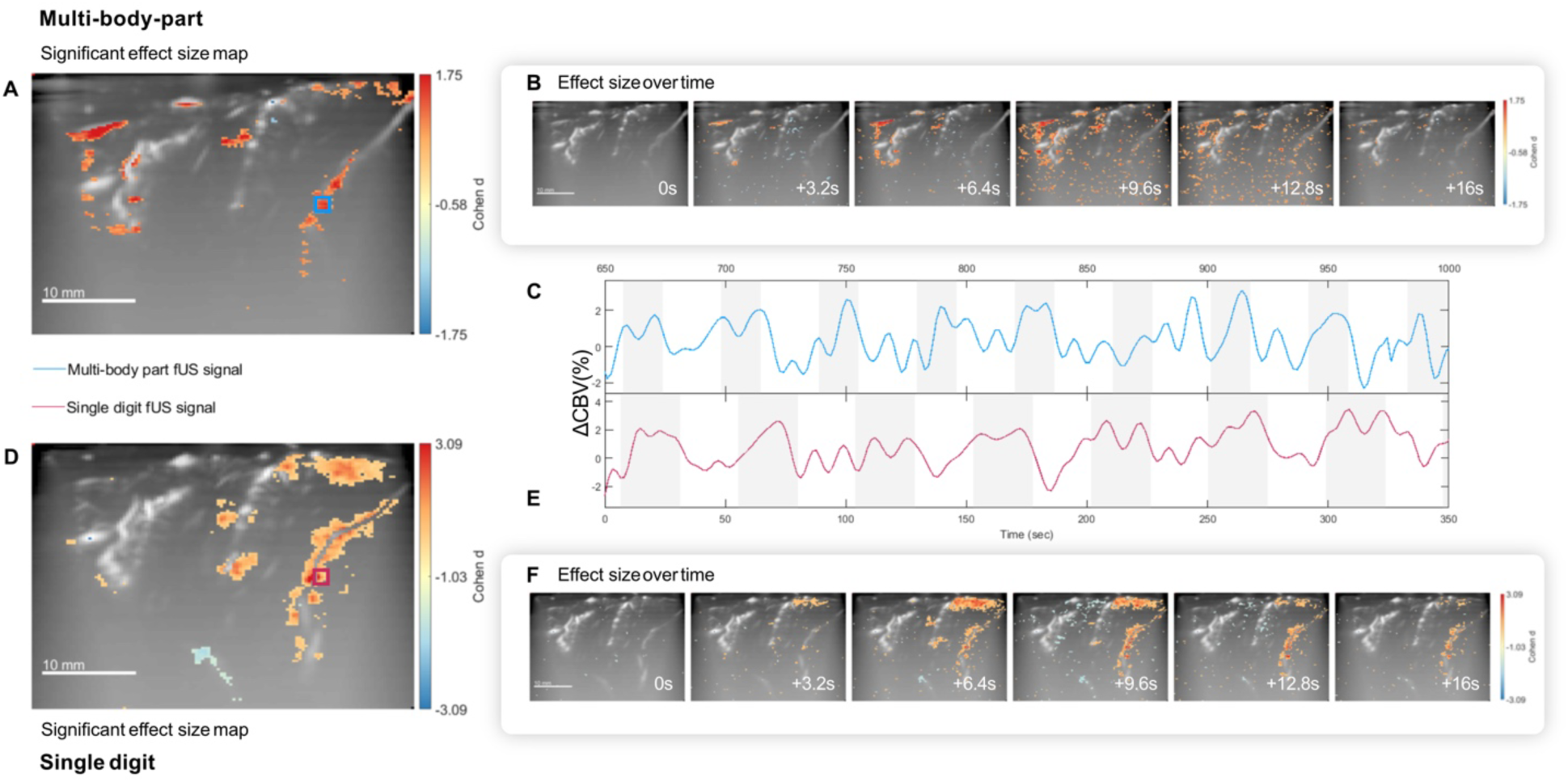
fUSI can detect single-trial event-related neurovascular activity. A-C) Multi-body-part movement. D-F) Single digit movement. A,D) Statistical parametric map of overall task activity and their corresponding Cohen d effect sizes (bonferroni correction, p<0.01). B,F) Statistical maps across different time points during execution for an arbitrarily selected effector – lip and middle finger, respectively. Significant event-related activity began at approximately 3.2 s after cue and peaked between 6-9 s for both multi-body-part and single digit movement (bonferroni correction, p<0.01). C,E) Average fUSI signal within the highlighted ROI (blue = multi-body-part, red = single digit) over time. Grey, shaded boxes indicate movement execution periods.

**Fig. 3:**
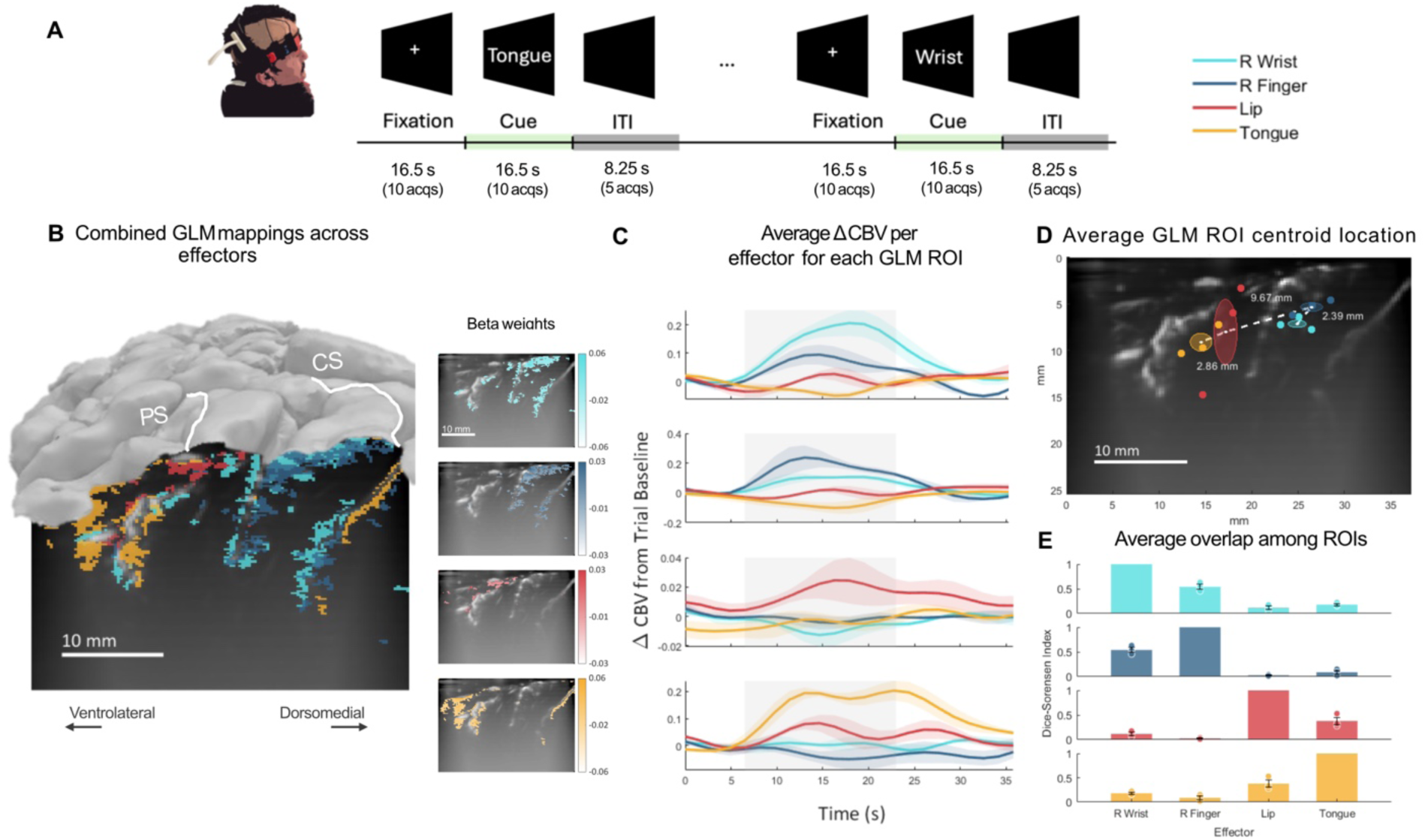
fUSI can robustly detect multi-body-part somatotopy in the sensorimotor cortex. A) Multi-body-part movement task design. B) Example single-session combined map of statistically significant ROIs generated from general linear modelling (GLM) for each effector – contralateral wrist, contralateral finger, lip, and tongue - and the individual ROIs per effector and their corresponding beta weights (p<1e-3 w FDR correction, white solid lines show the cortical surface and sulci, PS = Postcentral Sulcus, CS = Central Sulcus). C) Percent change in CBV per ROI for each effector represented as mean ± SEM, averaged across sessions. D) Centroids of activation per ROI averaged across sessions. Centroids were calculated based on the regions of activation per condition within the S1 subregion and are indicated by a colored dot corresponding to each effector per session. Average centroid position was displayed as ellipses with the mean as the center of the ellipse and ± SEM as the x and y dimensions of the ellipse. The white dotted lines between averaged centroids represent the connections between adjacent effectors based on the somatotopic model, with calculated Euclidian distances between centroids being displayed by their corresponding lines. E) Session averaged overlap among each ROI compared to the other ROIs using the Dice-Sørensen index. For each graph, one ROI, indicated by color, was used as a reference to calculate the Dice-Sørensen index compared to all the other conditions’ ROIs, shown as mean ± SEM. This represents how similar the activity distribution for a condition is to all the other conditions. Dice-Sørensen indices from individual sessions were reported as single dots on the bar graph.

**Fig. 4:**
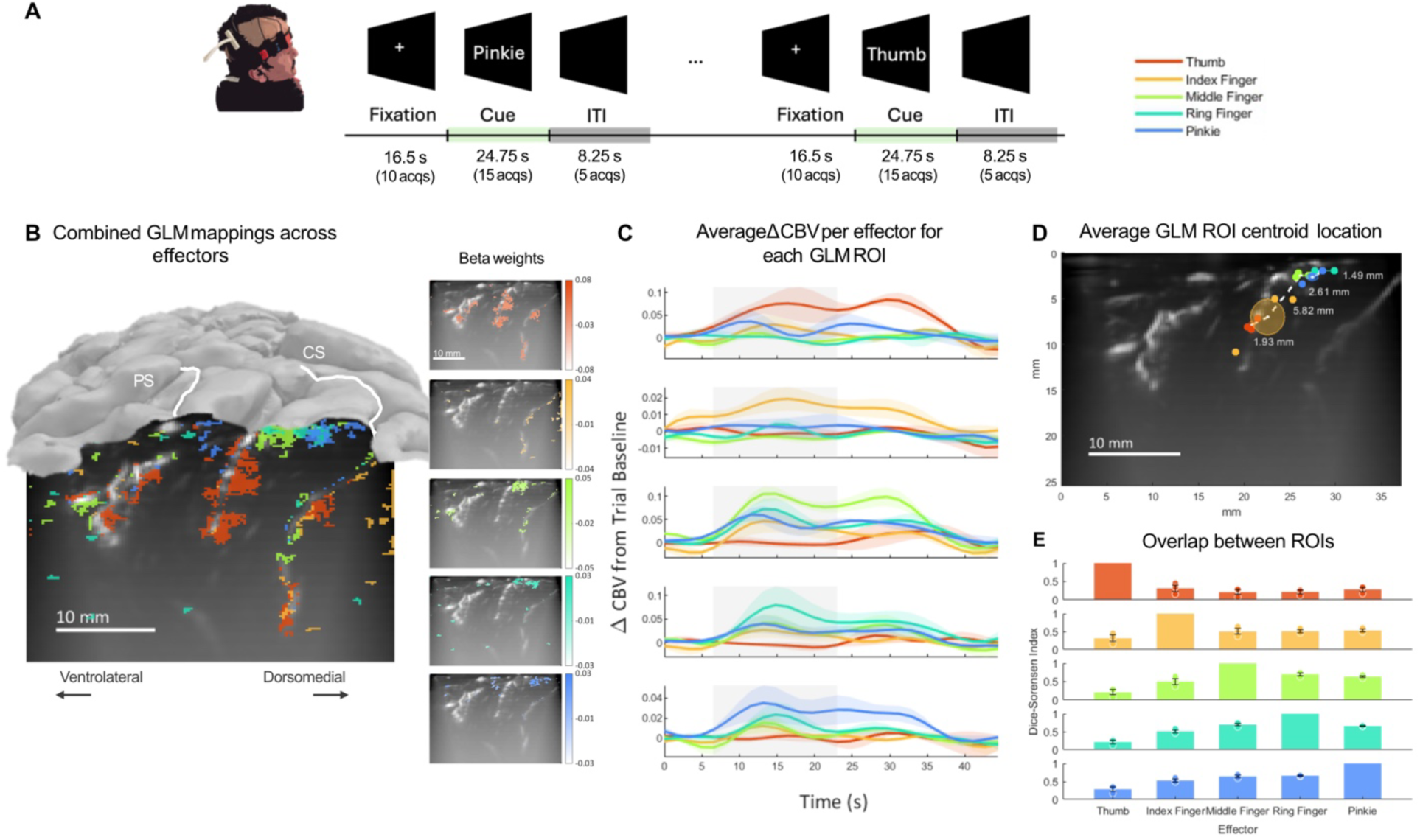
fUSI can identify single digit activity during individual finger movement in the sensorimotor cortex. A) Single digit movement task design. B) Example single-session combined map of statistically significant ROIs generated from GLMs for individual fingers and the ROIs per finger and their corresponding beta weights (p<1e-3 w FDR correction, white solid lines show the cortical surface and sulci, PS = Postcentral Sulcus, CS = Central Sulcus). C) Average percent change in CBV per ROI for each finger represented as mean ± SEM, averaged across sessions. D) Centroids of activation per ROI averaged across sessions. Centroids were calculated based on the regions of activation per condition within the BA 1 subregion and are indicated by a colored dot corresponding to each effector per session. Average centroid position was displayed as ellipses with the mean as the center of the ellipsoid and ± SEM as the x and y dimensions of the ellipse. The white dotted lines between averaged centroids represent the connections between adjacent effectors based on the somatotopic model, with calculated Euclidian distances between centroids being displayed by their corresponding lines. E) Session averaged overlap among each ROI compared to the other ROIs using the Dice-Sørensen index. For each graph, one ROI, indicated by color, was used as a reference to calculate the Dice-Sørensen index compared to all the other conditions’ ROIs, shown as mean ± SEM. This represents how similar the activity distribution for a condition is to all the other conditions. Dice-Sørensen indices from individual sessions were reported as single dots on the bar graph.

**Fig. 5:**
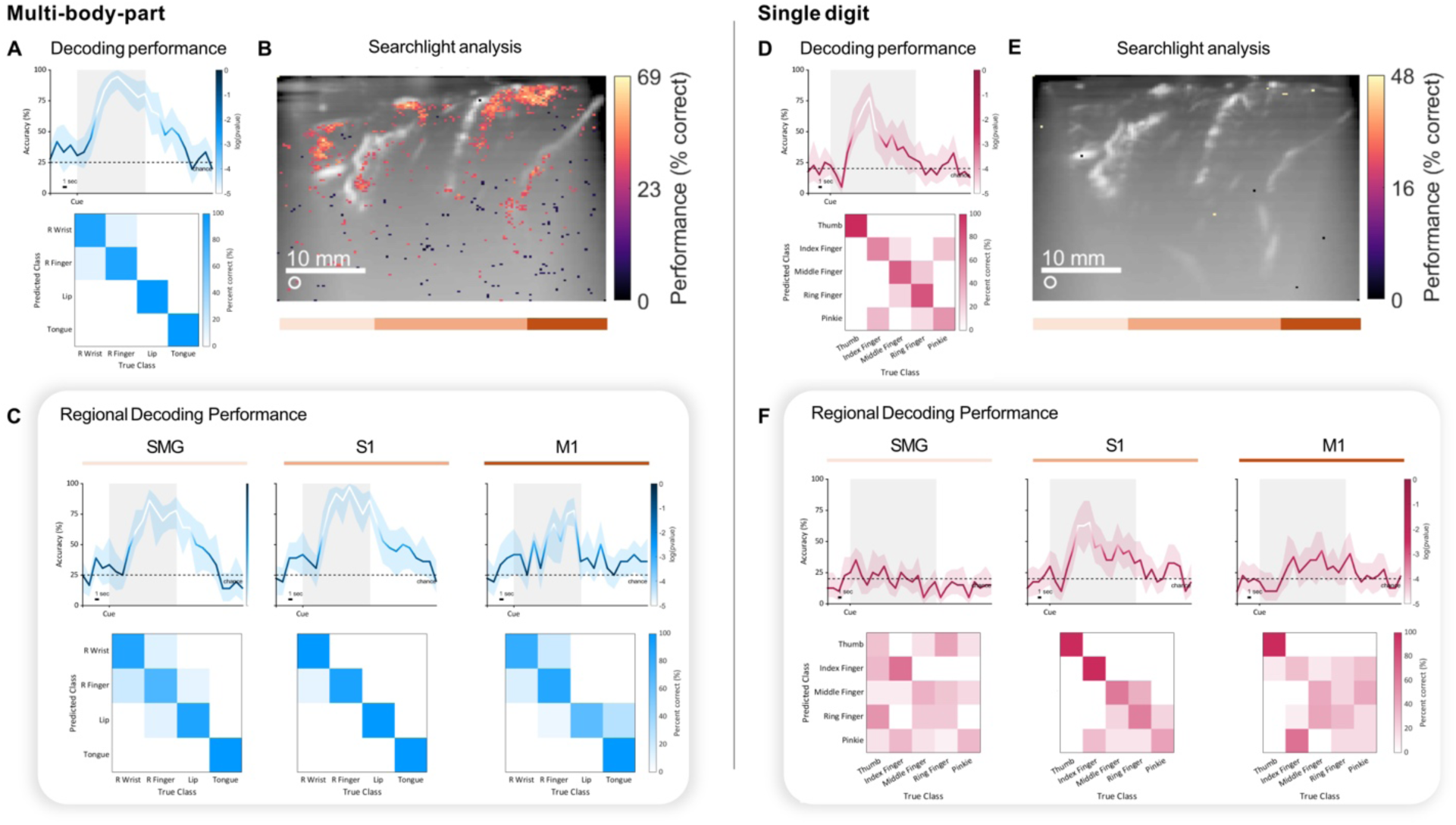
Multi-body-part and single digit movement effector information can be decoded from fUSI recordings of the sensorimotor cortex. A-C) Multi-body-part movement. D-F) Single digit movement. A,D) Movement effector decoding accuracy over time and the corresponding confusion matrix for the timepoint with the best decoding accuracy. Peak decoding reaches >90% accuracy for multi-body-part movements and >75% accuracy for single digit movements. The shaded region shows the execution period. The color of the decoding accuracy line shows statistical significance (1-sided binomial test). Confidence intervals were calculated using bootstrapping over 1000 permutations. B,E) Searchlight analysis of local regions containing sufficient information for decoding movement effector information (searchlight radius = 600 μm, FDR correction, p<0.05). The fUSI plane was then segmented into separate brain regions - SMG (peach), S1 (light orange), M1 (dark orange) - to examine the contributions of individual brain regions to decoding. C,F) Decoding accuracy over time and the corresponding confusion matrix for the timepoint with the best decoding accuracy using significant task-correlated voxels only within SMG, S1, and M1. The shaded region shows the execution period. The color of the decoding accuracy line shows statistical significance (1-sided binomial test).

**Fig. 6:**
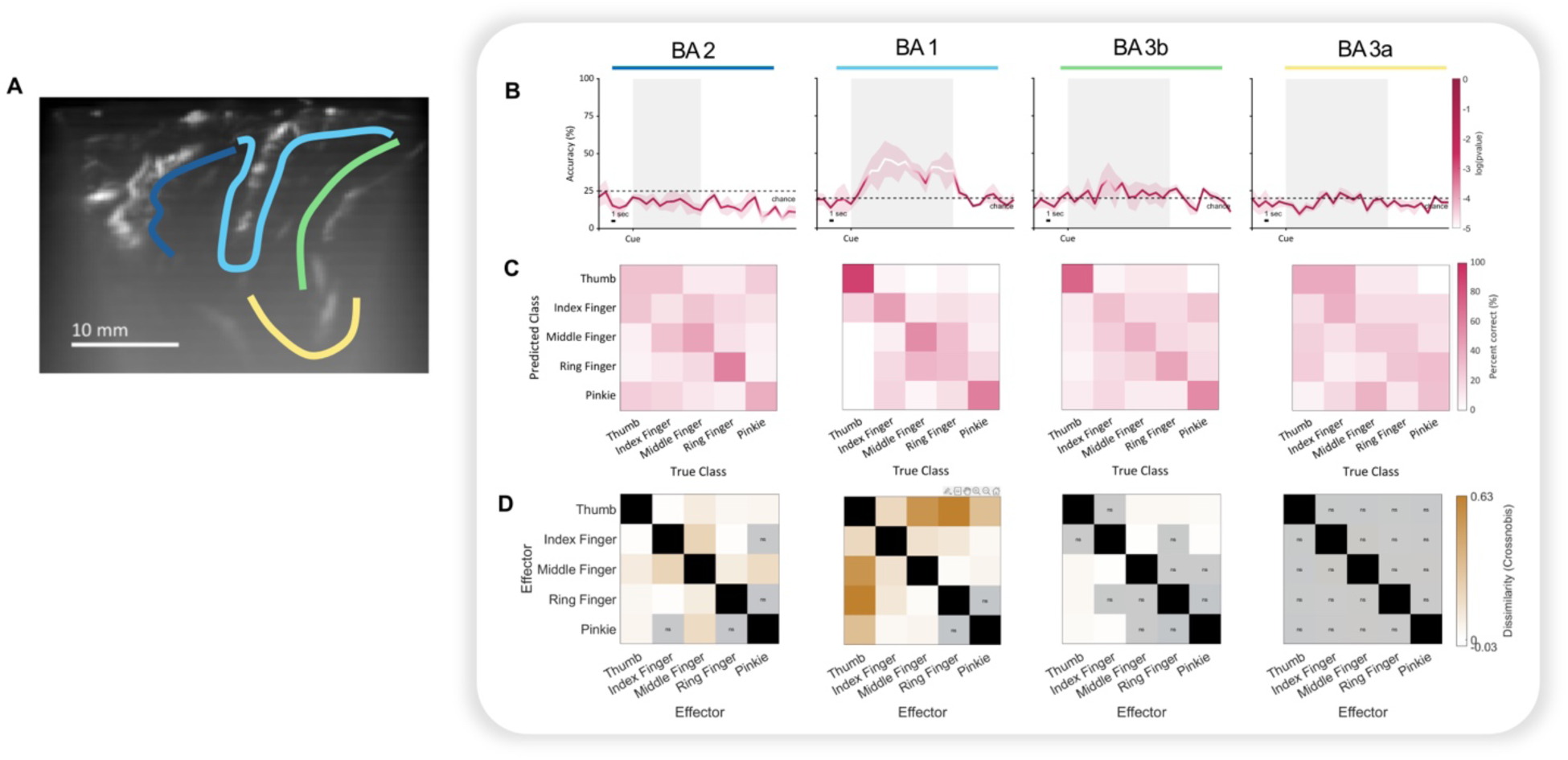
Motor effector representation varies across cytoarchitectonic regions. A) Segmentation of fUSI plane into BA 2 (dark blue), 1 (light blue), 3b (green), and 3a (yellow) based on probabilistic maps and anatomical definitions. Decoding and RSA was performed using the fUSI signal within the entirety of each segment. B) Average decoding accuracy over time. Only BA 1 produced significant decoding performance above chance. The shaded region shows the execution period. The plotted line shows mean ± SEM, n = 3 sessions. Color of the decoding accuracy line shows statistical significance (1-sided binomial test). C) The corresponding confusion matrix for the timepoint with the best decoding accuracy for each BA. D) RDMs generated for each BA to compare representational differences across cytoarchitectonic regions. RDMs were generated by calculating the Crossnobis distances from trial-averaged signal per voxel at peak activation for each effector and each run. Shaded boxes marked with the symbol “ns” represent nonsignificant Crossnobis values.

**Fig. 7:**
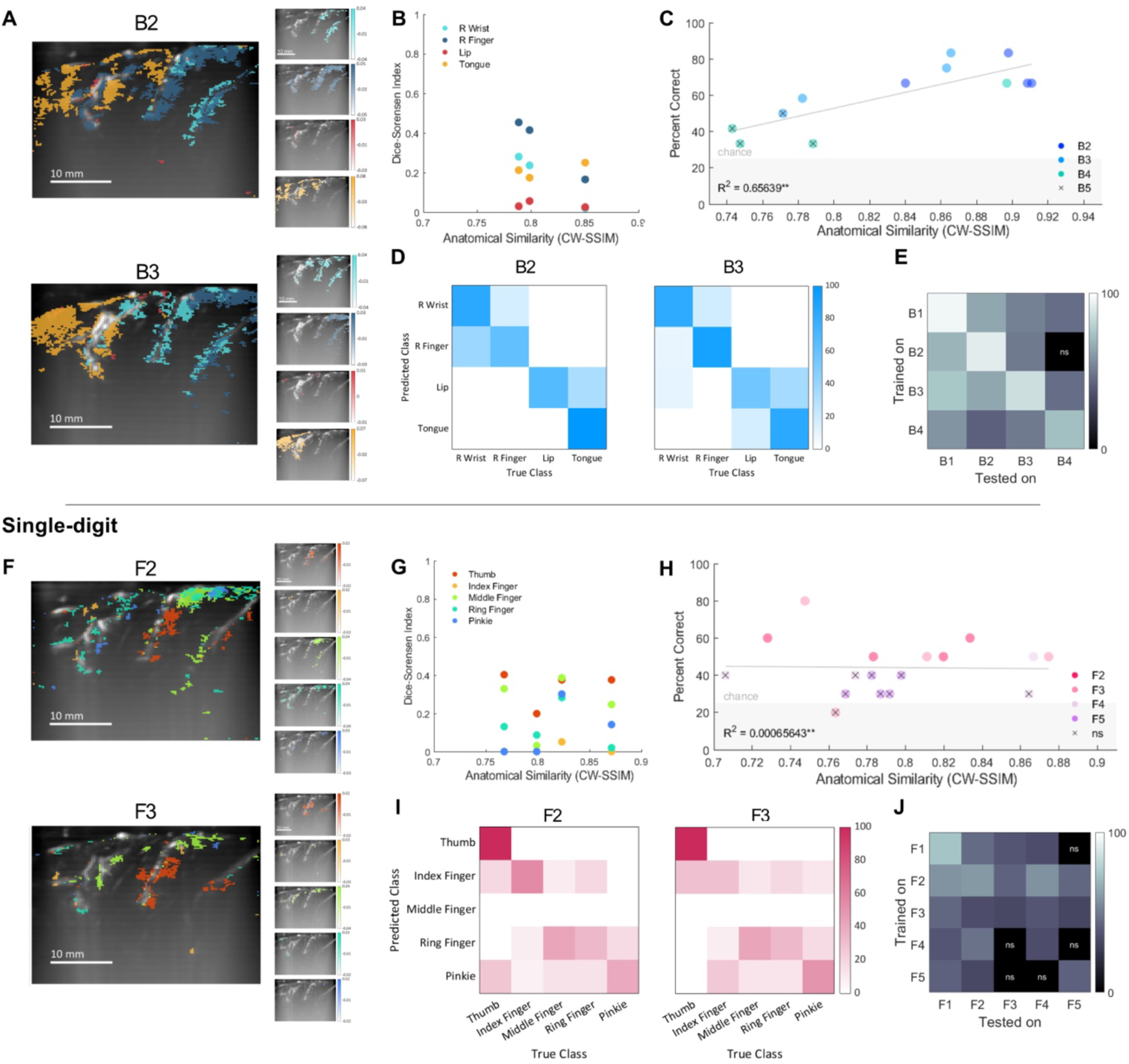
fUSI can provide stable mappings of similar neuronal populations across sessions. A-E) Multi-Body-part (B) movement. F-J) Single digit (F) movement. A,F) Comparison of GLM mappings generated from two separate sessions acquired on different days. B,G) The Dice-Sørensen indices representing the degree of overlap between each session’s GLM maps for individual effector activity relative to each session’s similarity in imaging plane position, represented by CW-SSIM. Each dot represents an individual effector for a given session. C,H) Across session decoding accuracy for individual trial runs (50-60 trials each) compared to each run’s anatomical similarity to the training session, session B1/F1, represented by CW-SSIM. Each dot represents an individual run, with runs being color coded by the session they came from. Grey line shows the linear trendline across all runs (correlation coefficient, **<0.01). Shaded region shows below chance level decoding accuracy. Nonsignificant decoding performance was indicated by an ‘x’ over the corresponding dot (1-sided binomial test). D,I) Example confusion matrices at the timepoints with the best decoding accuracies for across session decoding. Across session decoding was tested on 240 trials per session for multi-body-part movement and 200 trials per session for single digit movement. E,J) Cross-session decoding trained and tested on all session pairs. Blacked out boxes marked with the symbol “ns” represent session pairs with nonsignificant decoding (1-sided binomial test).

## Results

To detect hemodynamic task-correlated signal, we used a linear array ultrasound transducer (center frequency 7.5MHz, 60% bandwidth) to acquire fUSI images at 0.6 Hz from a human participant with an acoustically transparent PMMA skull implant while they performed cued movement tasks moving various body parts or “motor effectors.” During experiment sessions, the transducer was placed normal to the skull implant on the skin and held in place using an augmented helmet to return to a similar position during subsequent sessions. The transducer was positioned in a region approximately above the sensorimotor cortex using estimates from the participant’s MRI, providing a large field of view (38.4 mm width, 49.3 mm depth) including the primary motor cortex (M1), primary somatosensory cortex (S1), and supramarginal gyrus (SMG) at a 300 μm by 300 μm spatial resolution. The fUSI plane was then manually aligned to the participant’s MRI to better localize the plane’s position using vascular fiducial markers. Large vessels visualized in the fUSI plane were matched to the outline of sulci and gyri in the anatomical MRI (Fig. S1a,b). The chosen plane was also functionally verified through basic finger tapping and finger vibration tasks.

The primary sensorimotor cortex, consisting of M1 and S1, is well-known to be involved in movement and sensation across the body. Sensorimotor signals are encoded in a general somatotopic distribution within both M1 and S1, with single digit signals partially overlapping, particularly within M1^24–27^. SMG, part of the posterior parietal cortex, is involved in sensory integration and movement planning, including for complex tool use^28^. This makes the identified fUSI plane well suited for testing fUSI’s capabilities in distinguishing activity associated with different body part movements.

### fUSI can map multi-body-part somatotopy in the sensorimotor cortex

We used fUSI to record from our participant while they performed cued movement tasks using various motor effectors across the body– contralateral wrist, contralateral finger, lip, and tongue - repeatedly in a blocked task design (Fig. 3a). To validate the sensitivity of fUSI, we used effectors that are known to lie further apart on somatotopic mappings in the sensorimotor cortex as an initial test of fUSI’s ability to distinguish differential effector activity. To accumulate sufficient trials for decoding, we aligned and concatenated different trial runs within the same session. Additionally, large vascular landmarks were masked out of the image to ensure results were physiologically relevant to brain parenchyma. Using fUSI, we were able to detect single-trial changes in neurovascular signal, represented by percent change in CBV, corresponding to task execution (Fig. 2c). To verify consistent event-related activity and identify significant task-correlated voxels, we generated statistical parametric maps of overall task activity using voxel-wise Student’s t-tests (two-sided with Bonferroni correction) between execution and fixation periods and calculated the Cohen’s d effect size of significant voxels (Fig. 2a). We used the same analysis to generate statistical maps across different time points during execution for an arbitrarily selected effector to understand spatiotemporal signal dynamics. Significant event-related activity began at approximately 3.2 s after cue and peaked between 6 - 9 s, highlighting voxels in SMG and S1 (Fig. 2b). Similar activation patterns were found for the other effectors. This matches expected activation patterns given the hemodynamic origins of the fUSI signal. We then used this information to generate fine-grained maps of voxels correlated with each movement effector.

Using general linear modeling (GLM) analysis, we generated T-contrasts for each effector relative to rest and identified statistically significant regions of interest (ROIs) linked to each effector across three different multi-Body-part (B) movement sessions: B1, B2, and B3. These ROIs indicated a dorsomedial to ventrolateral distribution for finger, wrist, lip, and tongue, respectively. To better visualize the distinct distributions of each effector, we generated T-contrasts comparing effectors against each other and used this for our mapping visualization and signal calculations. We show the T contrasts from session B1 as an example (Fig. 3b). When examining the average neural activity across all conditions for each ROI, we found that average neural activity for a given ROI was highest for its corresponding condition (Fig. 3c). To compare the distributions of each effector, we calculated the centroids, weighted by activity, of the ROIs found for each condition across sessions and compared their positions. Only ROIs within S1 were used to calculate the centroids to better capture somatotopy. Centroid locations followed general canonical somatotopic mappings of movement effectors in S1^24,29^ (Fig. 3d). From dorsomedial to ventrolateral, average centroid separation increased from wrist to finger (2.39 mm), finger to lip (9.67 mm), and lip to tongue (2.86 mm), displaying the classic somatotopic pattern of same limb effectors lying closer together and cross body effectors lying further apart. Additionally, we calculated the average amount of overlap between effector representations across sessions using the Dice-Sørensen index, which compares the number of intersecting voxels between representations to the sum of voxels in each representation. The Dice-Sørensen indices were calculated for each pairwise combination of effectors, generating a matrix, and then averaged across sessions. The average Dice-Sørensen indices showed that finger and wrist display a large proportion of overlap (0.54) with little to no overlap with lip and tongue (0.12-0.18). Lip and tongue similarly had a large degree of overlap between themselves (0.38), though to a lesser extent than finger and wrist (Fig. 3e). This further aligns with the traditional somatotopic homunculus, in which finger and wrist representation lie closer to each other and lie further from lip and tongue representation.

### fUSI can map individual finger movement in the sensorimotor cortex

Having established that fUSI can be used to distinguish movements across different body parts, we conducted additional experiments on distinguishing individual finger representation in the sensorimotor cortex, which requires more refined spatial resolution given the mixed representation of intralimb effectors. We used the same cued movement task format as the prior experiment but, this time, asked the participant to move individual fingers on their contralateral hand (Fig. 4a). Applying the same analysis as the prior experiment, we were able to detect task-correlated hemodynamic changes in signal (Fig. 2e). Statistical parametric maps of general task activity showed activation within the dorsomedial region of S1 (Fig. 2d). When examining changes in significant task-correlated voxels over time during execution for the middle finger, significant event-related activity began around 3.2 s and peaked between 6-9 s, similar to that of multi-body-part movement (Fig. 2f). Of note, voxels within BA 1 were activated first, with voxels within BA 3b and 4 activating shortly afterward and persisting after BA 1 activity diminished. This pattern was seen for the other fingers as well.

We then generated fine-grained GLM mappings and were similarly able to identify ROIs tuned to each individual finger across multiple individual Finger (F) movement sessions: F1, F2, and F3. These maps exhibited a clear dorsomedial to ventrolateral distribution for pinkie to thumb activity, respectively, within the sensorimotor cortex as shown by session F1’s mappings (Fig. 4b). This is shown by the shift in average centroid location from dorsomedial to ventrolateral for pinkie to thumb representations, with the centroids for pinkie and ring finger lying 1.49 mm apart, ring to middle lying 2.61 mm apart, middle to index lying 5.82 mm apart, and index to thumb lying 1.93 mm apart (Fig. 4d). Centroids for single digit representation were calculated using ROIs within BA 1 to more clearly visualize somatotopy. Average neural activity across sessions per ROI confirmed that activity was highest for the corresponding finger, though the ROIs for the middle, ring, and pinkie fingers exhibited more mixed activity across the various fingers (Fig. 4c). The average Dice-Sørensen indices across sessions also were higher for ROIs of neighboring fingers, especially for the middle and ring fingers, suggesting a subtly shifting distribution for individual finger representation with more overlap between the middle and ring fingers. Dice-Sørensen indices were on average higher (0.57) compared to the values seen in multi-body-part movement (0.42), supporting prior findings of intermixed encoding for single digit movement within the sensorimotor cortex^25–27^. Dice-Sørensen indices ranged from 0.21-0.70 (Fig. 4e).

### Multi-body-part and single digit movement effector information can be decoded from fUSI recordings of the sensorimotor cortex

Given our success in mapping fine-grained body representations with fUSI, we then examined whether fUSI data captured sufficient information for decoding single-trial motor effector information. We first generated statistical maps of overall task activity using only the first run of a given session, which was then excluded from decoding analysis. These statistical maps were used as binary masks to exclude task-irrelevant voxels during decoding. Principal component analysis (PCA) was used to reduce the dimensionality of the remaining input data - masked, flattened fUSI images - and then linear discriminant analysis (LDA) was applied to classify the resulting principal components. All reported results were generated using 10-fold cross-validation. Using this method, we were able to achieve significant decoding above chance level for multi-body-part motor effectors at >95% within 36 trials in session B1. Individual finger movements could also be decoded at above chance level at >75% within 40 trials in session F1, with errors primarily occurring between neighboring digits (Fig. 5a,d).

To identify features important for distinguishing motor effectors during decoding, we mapped out the top 1% highest weighted voxels from a PCA-LDA classifier trained on an entire fUSI image acquired at the timepoint that produced peak decoding. Voxels were identified by taking the LDA weights from the PCA-LDA decoder, which represent PCA features that best discriminate between different motor effector pairs, and projecting the features to voxel-space using the inverse PCA transform. For multi-body-part movements, decoding weight maps showed that the highest weighted voxels tended to cluster in small regions of SMG, S1, and M1, specifically in BA 1, 3b, and 4 (Fig. S4a). For single digit movements, the weight maps concentrated mainly in superficial layers of the sensorimotor cortex, clustering at BA 1 and 3b (Fig. S4b). The highest weighted voxels from the PCA+LDA decoder lie within the statistical parametric maps and GLM mappings of general task activity, indicating that voxels used the most for decoding align with functionally relevant regions of the brain rather than noise regions or other confounds.

We also wanted to examine whether local regions contained sufficient information for high accuracy decoding. To do so, we performed a searchlight analysis with a 600 μm radius. Analyses depicted clusters of voxels with high decoding accuracy within the sensorimotor cortex and SMG for multi-body-part movement, indicating that local neighborhoods across SMG, S1, and M1 contain sufficient information for decoding multiple effectors across the body (Fig. 5b). Single digit movement similarly contained voxels with high decoding accuracy within the sensorimotor cortex, though the searchlight analysis only exhibited a few individual voxels involved in decoding rather than the large clusters seen for multi-body-part movements (Fig. 5e). This may reflect the more distributed and overlapping nature of single digit representations, resulting in noisier signal in local regions and requiring integration of information across broader spatial regions to have significant decoding performance.

Seeing how features and voxels containing decodable information were interspersed across various brain regions, we wanted to examine how decoding performance varied across these individual regions. The fUSI plane was segmented into SMG, S1, and M1, and the significant task-correlated voxels within each of those ROIs was used for decoding.

For multi-body-part movement, S1 produced the highest peak decoding accuracy of 97%, following the well-defined somatotopy seen in S1. M1, in comparison, had the lowest peak decoding accuracy of 76%, which reflects the intermixed somatotopy seen in M1. Most errors in both S1 and M1 were between adjacent motor effectors, finger and wrist vs. lip and tongue, further supporting the presence of somatotopic distributions. Interestingly, SMG had robust decoding performance at 87%, indicating that SMG contains sufficient information for decoding movement parameters and providing a potential target for decoding movement in the future (Fig. 5c).

For single digit movement, S1 also produced the highest peak decoding accuracy at 64% while M1 had a peak decoding accuracy of 56%. SMG did not display significant decoding performance (Fig. 5f). This aligns with the expected somatotopic distributions seen in S1 and M1, especially given that single digit movement distributions have been shown to be highly overlapping. The lack of significant decoding performance in SMG compared to multi-body-part decoding is possibly due to insufficient local information within the small field of view for SMG. This is emphasized by the small number of voxels found to be important for decoding in SMG in PCA-LDA weight maps and during searchlight analysis.

### Movement effector information is variably encoded across different cytoarchitectonic regions of the sensorimotor cortex

Given that voxels important for decoding clustered in specific subregions of S1, we wanted to go further and determine whether movement effector information was encoded differently across various cytoarchitectonic regions. To do so, we examined the decoding performance and representational similarity across anatomically defined regions corresponding to BA 1, 2, 3a, and 3b for single digit movement. We first segmented S1 into its corresponding BAs by combining probabilistic maps and anatomical descriptions^30–32^ of BAs (Fig. 6a). Given that our participant had an anatomical variation resulting in a sulcus running through the crown of S1, we defined the sulcus lying between the central sulcus and postcentral sulcus as part of the crown of S1 and, in turn, part of BA 1. We then used the entirety of each segmented BA to perform decoding and generate representational dissimilarity matrices (RDM)^33^. To generate RDMs, we computed pairwise Crossnobis distance — a noise-normalized, cross-validated measure of representational dissimilarity — for each finger pair, averaged across sessions (see Methods). Distances are expected to be zero under the null hypothesis given that two patterns are identical with respect to the noise level. Positive distances indicate patterns are more dissimilar to each other. Negative distances can also occur but are typically caused by incidental anticorrelation in the setting of noise and variability across sessions. Interestingly, results demonstrated that only BA1 contained sufficient information for significant decoding at a peak decoding accuracy of 48%. All other BAs did not produce significant decoding performance, though BA3b had a slight increase in decoding accuracy (Fig. 6b,c). RDMs revealed that BA 1 exhibited the largest dissimilarities between fingers, with Crossnobis values reaching a peak of 0.63 while BA 2, 3b, and 3a had maximum values of 0.21, 0.04, and 0.02, respectively (Fig. 6d, S5). These larger distances between fingers may explain the significant decoding accuracy when using only BA 1. These results are unexpected given that BA 3b has previously been reported to be involved in low-level cutaneous tactile input processing, having small receptive fields and strong somatotopy^34^. In comparison, BA 1 and BA 2 receive inputs from BA 3b and demonstrate hierarchical, higher-level integration of tactile information. These findings may be a result of the limited plane of view and position of the plane, and further study is necessary to better understand these cytoarchitectural differences. Overall, the index, middle, ring, and pinkie fingers exhibited small distances across the BAs consistent with the overlapping finger representations commonly seen for intralimb effectors. In comparison, the thumb exhibited larger distances from all other fingers, a similar pattern seen in naturalistic hand use models of activity^35^.

### fUSI can provide stable mappings of similar neuronal populations that can be used for cross-session decoding

One of fUSI’s strengths is that it provides reproducible imaging of similar neuronal populations^19^. This allows us to reliably examine behavioral representations from the same neuronal populations across different sessions. We demonstrate this by comparing the prior generated GLM mappings for multi-body-part and single digit movement from sessions B1 and F1, respectively, to additional sessions recorded on different days. To compare the overlap between regions of activity identified by GLM, we calculated the Dice-Sørensen index for individual effectors. For multi-body-part movement, session B2 and B3 exhibited effector representations qualitatively similar to those of session B1 (Fig. 7a). However, the Dice-Sørensen index averaged across effectors was 0.24 for session B2, 0.22 for session B3, and 0.12 for session B4, resulting in an overall average 0.19 Dice-Sørensen index across all effectors and sessions (Fig. 7b). For single digit movement, sessions F2 and F3 displayed similar patterns in effector distribution to those of session F1, with finger representation moving dorsomedial to ventrolateral for pinkie to thumb ROIs, but also had clear differences in each individual effector’s representation (Fig. 7f). The Dice-Sørensen index was 0.28 for session F2, 0.16 for session F3, 0.17 for session F4, and 0.1 for session F5, for thumb to pinkie, respectively (Fig. 7g). The average Dice-Sørensen index across all fingers and sessions was 0.17. This indicates that while representations across sessions follow similar patterns, individual representations per session may vary across days. This could be due to variations in experimental setup across sessions such as deviations in plane position.

To determine the influence of plane position on the reproducibility of effector mappings, we calculated the complex-wavelet structural similarity index measure (CW-SSIM)^19,36^ of each session relative to sessions B1 and F1, respectively, as a representation of imaging plane anatomical similarity across sessions. Using this measure, there was no clear relationship between plane similarity and Dice-Sørensen overlap scores for both multi-body-part and single digit movement (Fig. 7b,g). The lack of relationship and low Dice-Sørensen scores may be because individual effector encodings are sparsely represented, particularly for single digit movement, resulting in poor overlap over minor differences across sessions. This is seen by how effectors with smaller ROIs, such as the lip, pinkie, and index finger, generally had lower Dice-Sørensen indices across sessions.

Given the similarity in the general distribution of effectors, we attempted to decode movement effector information from one session using a PCA+LDA decoder trained on another session. Since the decoder trained on sessions B1 and F1 had the best performance out of all the sessions for multi-body-part and single digit movement respectively, we used those sessions’ decoders to decode the remaining sessions without additional training. For multi-body-part movement, we were able to reach a peak decoding accuracy of 70% and 72% for sessions B2 and B3, with most errors being made for wrist vs. finger and lip vs. tongue (Fig. 7d). For single digit movement, we reached a peak decoding accuracy of 45% and 40% for sessions F2 and F3 (Fig. 7i). Interestingly, both sessions F2 and F3 failed to predict the middle finger during decoding. This could be due to the increased overlap between finger representations, especially for the middle finger, making the decision boundary more vulnerable to minor shifts in imaging plane across sessions. Given this performance, we then tested cross-session decoding across all pairs of sessions to demonstrate that effector representations remain robust regardless of session. For multi-body-part movement, cross decoding maintained above chance performance for all session pairs except for those involving session B4, which also had the lowest Dice-Sørensen overlap scores (Fig. 7e). For single digit movement, cross decoding performed significantly above chance for most session pairs (Fig. 7j). These findings demonstrate that while individual effector representations may vary across sessions, general patterns remain robust regardless of session and can be used for decoding across sessions.

Lastly, to understand the sensitivity of cross-session decoding to shifts in imaging plane position, we compared the decoding accuracies of individual runs to their CW-SSIM values relative to sessions B1 and F1. We used a decoder trained on sessions B1 and F1 for multi-body-part and single digit movement, respectively. We found that decoding accuracy had a positive correlation with CW-SSIM for multi-body-part movement and little to no correlation with CW-SSIM for single digit movement (Fig. 7c,h). This demonstrates that decoding using fUSI is tolerant to plane shifts for coarse representations such as multi-body-part representations where the decoder can generalize better but less so for fine-grained, overlapping representations such as those seen for individual fingers, requiring more refined, precise positioning.

## Discussion

This work demonstrates that fUSI is a robust neuroimaging tool that can be used to accurately map and decode motor effector information from neural activity in a human participant through an acoustically transparent cranial window. By mapping and decoding both multi-body-part and single digit movement effectors from fUSI signals, we show that fUSI can robustly access rich and spatially organized information within the sensorimotor cortex across different experiment sessions. These results build upon prior NHP^17–19^ and human work^13,20,37^ on mapping and decoding with fUSI.

The ability of fUSI to resolve differential motor effector organization across the sensorimotor cortex demonstrates its potential as a mesoscopic imaging modality capable of bridging the gap between population-level hemodynamics and cellular electrophysiology. Prior studies have already used fUSI for mesoscopic mapping in various organisms including olfaction in rats^38^, auditory stimuli in ferrets^39^, movement planning in NHPs^19^, and functional and vascular brain maps in intraoperative human patients^40^. Our multi-body-part mappings reproduced the canonical dorsomedial-to-ventrolateral organization of sensorimotor effectors seen in classic Penfield mappings and modern fMRI results^24,27^. More importantly, fUSI was able to capture fine-grained single digit organization, distinguishing between individual finger activity within overlapping cortical regions and differences in encoding across cytoarchitectonic regions. Prior studies using electrophysiology have identified intermixed representation for individual finger movements in the sensorimotor cortex^26^. Previous fMRI studies have similarly identified overlapping distributions for individual finger representation, though they have also identified subtle somatotopic organization at the population level when comparing isolated finger representations^25,41^. fUSI is capable of capturing a clear dorsomedial-to-ventrolateral shift in overlapping representation for pinkie to thumb signal, respectively, that is consistent with these studies. The fact that these fine distinctions can be visualized and decoded from a single fUSI plane through skin and an acoustically transparent skull implant underscores the high spatial resolution and sensitivity of fUSI and vouches for its research potential.

It is important to note that these results are a case study of a single participant. Findings seen in the current participant may differ in other participants, and, thus, future studies are necessary to explore fUSI signal in additional participants. Additionally, since we are using a linear array probe, we are limited to one 2D imaging plane at a time. As a result, our mappings and analysis of multi-body-part and single digit movement may not be representative of larger-scale patterns that could be more apparent from a 3D view of the brain. While we can decode at significantly higher than chance from our current fUSI frame, volumetric fUSI is expected to further improve decoding abilities by providing increased information density and spatial information for mapping and image registration. Advances in volumetric fUSI have already emerged such as through matrix array^42^ and row-column array transducers^43^. Future studies will aim to apply such volumetric probes to further improve fUSI sensitivity and decoding abilities.

Using fUSI, we were able to decode multi-body-part movement at a peak accuracy of 95% and single digit finger movement at a peak accuracy of 78%, out-performing other minimally invasive neuroimaging techniques. Using fMRI, researchers decoded single touch signals from the left and right hand and feet at a peak accuracy of 70%^44^ and single movement signals from individual fingers at a peak accuracy of 63%^45^. This further exhibits the high sensitivity and spatial resolution at which fUSI is able to capture task-correlated activity, allowing for more accurate decoding. Coupled with fUSI’s ability to acquire neurovascular data epidurally and reliably record from similar neuronal populations long-term, these findings position fUSI as a contender for more accessible, minimally invasive BCI alternatives in humans, a point of active research.

Our decoding weight maps localized to cortical regions within M1, S1, and SMG—areas known to contribute to movement execution and somatosensory feedback. This shows that our decoder is drawing from information from areas known to be functionally relevant for our task rather than overfitting or decoding from confounding information such as movement artifact. Interestingly, the searchlight analysis for individual finger movements identified only a few scattered voxels capable of significant decoding using local information while LDA weight maps highlighted several clustered regions within the somatosensory cortex. This discrepancy between searchlight and LDA weight maps may suggest that single digit representations are encoded in globally distributed patterns rather than local neighborhoods at the population level.

A caveat of the current study is that tasks had to be performed in a blocked task format to generate sufficient activation for fUSI detection, likely due to signal attenuation from the PMMA implant. PMMA was found to result in reduced intensity and SNR^13^, and studies in which the skull bone was removed and the brain was directly imaged with fUSI report the ability to detect individual movements for reaches, saccades, and individual finger movements^17–19,37^. However, the PMMA implant is necessary to safely implement chronic fUSI recording in patients and has been shown to have improved SNR values compared to imaging through skull bone and titanium mesh, a common skull bone replacement. Increasing SNR by adjusting fUSI parameters such as voltage and half-cycle offers a potential solution, but these changes also introduce increases in temperature and mechanical pressure, which are essential to be within FDA limits for participant safety. Thus, careful tuning of fUSI parameters will be necessary to address this challenge. Implanted fUSI transducers may also offer an alternative for longitudinal recording of high-quality neurovascular signal. However, implanted transducers are still in an early stage of development^37,46^ and require extensive future exploration before considering implantation in a human subject.

Additionally, fUSI is limited by the hemodynamic nature of its signal, making it unable to match the temporal resolution of electrophysiology for decoding. Instead, fUSI’s strengths are best suited for slow-moving cognitive states, such as mood and movement intention, allowing for chronic monitoring in more naturalistic settings. For instance, because fUSI transducers are smaller and more compact compared to large fMRI, PET, and CT scan machines, they could be adapted for bedside monitoring, home neurorehabilitation, or outpatient BCI use. Research has already established precedents for clinical fUSI use, including bedside functional brain monitoring in neonatal patients^47^ and intraoperative brain mapping during tumor resection^40^. As noted above, fUSI systems are still restrained by the large computational systems required to store and process fUSI data, limiting motion to seated actions, actions within a certain radius, or actions coupled with movable fUSI systems. However, several potential solutions are already being developed such as conformal ultrasound transducer patches^48^. Methods to reduce the size of ultrasound computational systems or omit the need for transducers to be tethered to large computational systems through wireless methods are also currently being developed. Together, these advantages suggest a new paradigm in which chronic ultrasound imaging could provide both scientific and clinical insight into human brain function.

In summary, the contributions presented here establish one of the first published demonstrations, to our knowledge, of the use of fUSI in a human for mapping and decoding multiple forms of movement through an acoustically transparent cranial implant. Our work builds on prior fUSI studies that laid the foundations for the development of fUSI and fUSI-BCI, and we hope to contribute to the growing body of this research by showing that fUSI can resolve single-trial multi-effector human motor behavior. This work acts as a proof of concept of fUSI’s abilities and is a stepping stone for further development of fUSI in broader applications, from BCIs to eventual clinical use, including long-term neuropsychiatric monitoring, exploring speech semantic representation, decoding movement intention and more.

## Methods

### Study design and subject details

All procedures were approved by the Institutional Review Boards of the University of Southern California (USC), Caltech (IR24-0204), and Rancho Los Amigos National Rehabilitation Hospital (RLA). The participant was recruited for a prior study^13^ and was fitted with a custom polymethylmethacrylate (PMMA) acoustically transparent cranial implant (Longeviti ClearFit) after a TBI requiring decompressive hemicraniectomy and cranioplasty of the left hemisphere. The custom PMMA implant has a thickness of 4 mm to match the participant’s nominal bone thickness and includes a 2 mm thinned window above the sensorimotor cortex, identified through fMRI during a finger tapping task, for improved SNR. The participant underwent MRI and fUSI imaging before and after he was fitted with the custom cranial implant with as many acquisitions as allowed by the authorized protocol. During the fUSI recording, the participant engaged in cued movement tasks in which they were cued to move a specific body part repeatedly for a block of time. All fUSI study sessions took place at Caltech, and all CT and MRI scans occurred at the Keck Hospital of USC.

### Behavioral setup

The participant was seated in a reclining chair with a 27-inch fronto-parallel screen (Acer XB271HU) positioned 70 cm in front of him. For initial brain mapping and offline decoding, visual stimuli were presented using a custom Matlab script using the Psychophysics Toolbox extension ^49–51^, which also outputted timing information that was stored for offline access. Data analysis was performed in MATLAB 2023a (MathWorks, Natick, MA, USA) using standard desktop computers.

### Behavioral tasks

The participant performed cued movement tasks using several different motor effectors in a blocked task design. Contralateral wrist, contralateral finger, lip, and tongue were used for examining multi-body-part motor effectors while contralateral individual fingers (thumb to pinkie) were used for examining single digit motor effectors. For initial brain mapping and offline decoding tasks, the trial began with fixation on a center cue for 16.25 s (10 fUSI acquisitions). A text cue describing what motor effector to use appeared for 16.25 s (10 acquisitions) for multi-body-part movement and 24.75 s (15 fUSI acquisitions) for single digit movement, during which the participant began executing the cued movement repeatedly until the text cue disappeared. There was an 8.25 s (5 acquisitions) intertrial interval (ITI) before the next trial began (Fig. 2A, 3A).

### fUSI sequence and recording

During each fUSI recording session, we placed the ultrasound transducer (128-element linear array probe, 7.5-MHz center frequency, 60% bandwidth, 0.3-mm pitch, Vermon, France) on the skin using ultrasound gel as a coupling agent. We reliably positioned the ultrasound transducer across recording sessions through visual alignment with prior session images and manual adjustment using an augmented probe holder helmet. The imaging field of view was 38.4 mm wide and reached a depth of 49.3 mm, which allowed us to simultaneously capture multiple cortical regions, including parts of M1, S1, and SMG. The current plane was functionally verified through various somatomotor tasks and anatomically identified by manually aligning the fUSI plane to the participant’s MRI.

We used a programmable high-framerate ultrasound scanner (Vantage 256, Verasonics Kirkland, WA, USA) controlled by custom MATLAB fUSI acquisition scripts to drive the ultrasound transducer and collect pulse-echo radiofrequency data. A custom plane-wave imaging sequence was used at a pulse repetition frequency of 4000 Hz. Plane waves were transmitted at five tilted angles (−6°, −3°, 0°, 3°, 6°), in which 2 accumulations at each angle were used to generate a single compounded frame at a 400 Hz frame rate. Each power Doppler image was then generated every 1.65 s (0.6 Hz) from the accumulation of 300 compounded frames.

Each ensemble of 300 compounded frames underwent singular value decomposition (SVD) clutter filtering to separate tissue signal from blood signal before outputting a final power Doppler image. SVD upper and lower thresholds per power Doppler were adaptively calculated using the correlation matrix between spatial singular vectors^52^. In-phase and quadrature sampled data was outputted and stored to SVD process offline.

### fUSI anatomical alignment

The acquired fUSI plane was manually aligned to the participant’s T1w anatomical MRI to approximate anatomical data corresponding to different regions of the plane. Large vascular fiducials in the fUSI plane were aligned with the corresponding edges of sulci and gyri within the MRI. The anatomical MRI was segmented and labelled using the BrainSuiteAtlas1 in BrainSuite23a. These labels were used to delineate the fUSI plane into SMG, S1, and M1. Statistical Parametric Mapping (SPM25) was used to process and segment the participant’s T1w MRI into cytoarchitectonic probability maps. The MRI was warped into standard MNI space and then aligned with BA probabilistic maps^30–32^ using the SPM Anatomy Toolbox Version 3.0^53–55^ to identify ROIs for BA 1, 2, 3a, 3b, and 4 (4a and 4p combined). The fUSI plane was segmented into BAs using a combination of these ROIs and their anatomical definitions^30–32^. Based on the brain region labels generated in BrainSuite23a and anatomical definitions, the participant possesses an additional sulcus between the central sulcus and post central sulcus. Thus, this additional sulcus was included as part of the “crown” of S1 and was included in BA 1.

### Across-session alignment

To maintain participant attention, trial runs were shortened to no more than 10 minutes per run. Because of this, multiple runs needed to be aligned both within and between sessions in order to accumulate sufficient sample size for training. Using methods from a prior study^18^, fUSI imaging planes were spatially aligned using a custom semiautomated intensity-based rigid-body alignment script. The script used MATLAB’s ‘imregtform’ function to apply rigid body-alignment and transformation to desired fUSI image planes and provided a manual option to shift and rotate prior sessions to visually match runs together.

### Data preprocessing

Prior to offline analysis, a low-pass filter with a cutoff at 0.1 Hz was applied to remove temporal noise. For decoding analyses only, a Gaussian spatial filter (σ = 0.6370, FWHM = 450 μm) was additionally applied to reduce spatial noise and improve classifier generalization; all other analyses used unsmoothed data to preserve submillimeter spatial resolution. We then applied voxel-wise detrending, in which the best line of fit across a run was calculated using baseline signal during fixation periods and then subtracted from the given voxel’s signal, to address baseline drift. This also scaled signal to the baseline cerebral blood volume, generating more human interpretable values and making signal ranges more similar across runs.

### Quantification and Statistical Analysis

All analyses were performed using MATLAB 2023a. Unless otherwise stated, a significant difference was considered p < 0.05. All offline analysis excluded voxels above the brain surface. For initial activity mapping and offline decoding experiments, a 10 s delay was applied to each trial of sessions B1 and F1 for multi-body-part and single digit movement, respectively, shifting each trial forward by 10 s, to account for the hemodynamic delay and intertrial effects.

### Statistical parametric mapping

General task activation maps were generated for each experiment by comparing activity during the execution period, adjusted to account for hemodynamic delay, and the fixation period using a two-sided Student’s t-test for each voxel. Task activation maps over time were generated by performing voxelwise comparisons of average activity across trials for the selected condition at each time point to the fixation period, similarly using a two-sided Student’s t-test for each voxel. Bonferroni multiple comparisons correction was applied based on the number of voxels tested. The Cohen d effect size was calculated for significant voxels (small effect = 0.2, medium effect = 0.5, and large effect = 0.8)^56^. Significant voxels were visualized as a heatmap based on Cohen d values overlaid onto the fUSI anatomical plane. The generated maps of task significant voxels were used as binary masks during decoding to exclude task-irrelevant voxels.

### General Linear Modelling (GLM) mapping

GLMs were used to identify significant regions of interest relevant to each task condition, providing a map of activity in the given fUSI plane. GLM regressors were generated by convolving timepoints of interest including cue presentation per condition and fixation with a single gamma hemodynamic response function (HRF)^57^, with time constant (τ) = 0.7, time delay (δ) = 3 s, and n = 3 s. The GLM was then fit using the convolved regressors and the preprocessed fUSI signal from each voxel, generating β-values per condition per voxel. β-values represent how much a given regressor contributes to the acquired signal. Larger β-values indicate more involvement of a given regressor while smaller β-values indicate less involvement. Statistical significance of the β coefficients was determined using a T contrast with false discovery rate (FDR) correction (p < 0.001). Two T contrasts were used: 1) effectors relative to rest (Fig. S2) and 2) effectors compared to other effectors (i.e. Thumb – (Index + Middle + Ring + Pinkie)) (Fig. 3,4,7). To better depict the differences in individual effector representation, T-contrast 2 was used for visualizations and average signal change calculations. GLM results were visualized as heat maps for each β coefficient overlaid on the fUSI anatomical plane, with only significant voxels being kept. Voxel clusters under 4 pixels in size were excluded to improve visualizations. The cluster threshold was chosen through random permutation testing in which condition labels were randomized and the general task GLM was generated repeatedly to generate a distribution of cluster sizes. The cluster threshold was set as clusters larger than the 95^th^ percentile of cluster sizes (4 pixels). To better display differences in GLM maps across different conditions, significant voxels from each condition were overlaid together in different colors on the fUSI anatomical plane. This was done for the top 3 best quality sessions to demonstrate reproducibility.

To compare the different distributions among GLM maps, the centroid of activation for each GLM map per condition was calculated by calculating the center of mass of the ROI, weighted by activation, in Euclidian coordinates. This was repeated across 3 sessions, The average centroid location and standard error mean in the x and y axes across all 3 sessions was then computed. The Euclidian distances between neighboring conditions were calculated, with both the centroids and the distances being superimposed on an anatomical reference plane. We also calculated the Dice-Sørensen index, a measure of overlap ranging from 0 (no overlap) to 1 (complete overlap), between two GLM maps, A and B, to determine the degree of overlap between each condition’s map.

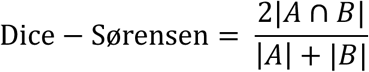

Dice-Sørensen values were displayed as a bar graph by calculating a matrix of Dice-Sørensen indices between each pair of effectors to demonstrate increased overlap of adjacent effectors. T-contrast 1 was used for Dice-Sørensen calculations to demonstrate the overlap of signal between effectors. This was repeated across 3 sessions, and then the session-averaged Dice-Sørensen calculations were reported.

### Image similarity

We calculated the pairwise complex-wavelet structural similarity index measure (CW-SSIM)^19,36^ of each session relative to sessions B1 and F1 for multi-body-part and single digit movement, respectively, as a representation of imaging plane similarity across sessions. The CW-SSIM measures the similarity of two images, where 0 is dissimilar and 1 is the same image. Sessions with higher CW-SSIM values were assumed to have less differences in imaging plane position while sessions with lower values were assumed to have larger shifts in imaging plane position. CW-SSIM is also more robust to small geometric distortions. We used an implementation freely available from the MATLAB Central File Exchange^58^ with 4 levels and 16 orientations.

### Session exclusion criteria

The pairwise CW-SSIM was calculated for all run pairs to determine the similarity of imaging planes across runs. Runs where CW-SSIM < 0.6 relative to runs from sessions B1 and F1 were deemed as different planes and excluded (Fig. S6). Sessions with fewer than 40 trials were also excluded from analysis due to insufficient data per session for decoding and mapping analysis.

### Decoding

Multi-body-part and single digit motor effectors were decoded at individual timepoints using principal component analysis (PCA, 95% variance kept) to reduce dimensionality and linear discriminant analysis (LDA) for classification. A PCA+LDA decoder is a commonly used method in prior fUSI decoding that is well-suited to classification problems with high dimensionality and small sample size. It was chosen since it was found to outperform other prior used decoders when using limited training sets. Training data was split into balanced sets for 10-fold cross-validation. Task irrelevant voxels were masked out of the dataset by applying a binary map of general task activity generated from the first run of a given session to improve decoding accuracy. Data used to generate these binary maps were excluded from decoding analysis to prevent circular analysis. Decoding accuracy was reported through percentage of correctly predicted trials and visualized through a single example session for both multi-body-part and single digit movement. Decoding performance for all other sessions is shown in Figure S3.

### Cross-session decoding

The same decoding pipeline as above was used to train a decoder on all time points of a given training session. The decoder with the best performance was then used to decode movement effector information across all time points for a different, unseen session. Cross-session decoding performance was reported through percentage of correctly predicted trials. To assess the performance of a decoder trained and tested on the same session, the 10-fold cross-validated percent accuracy was reported instead of the percent accuracy based on the entire test session.

### LDA weighting maps

LDA weighting maps were used to identify what voxels in the given anatomical plane were being used for decoding. To generate the maps, LDA weight coefficients across PCA components between each class were projected back into voxel-space. The weight maps were then overlaid onto the corresponding fUSI anatomical plane, keeping only the 1% highest weighted voxels.

### Searchlight analysis

Searchlight analysis was used to identify what voxels in the given anatomical plane were most important for decoding performance. It is a commonly used method in fMRI research that iteratively measures and maps the decoding performance within small regions of interest or “searchlights” centered on each voxel in the chosen image^59^. To perform searchlight analysis, we defined a circular region of interest (ROI; 600-μm radius) and performed offline decoding with 10-fold cross validation using a PCA + LDA classifier as described in prior studies^18^. This was repeated for each voxel in the imaging plane. To visualize the results, we overlaid the percent correct decoding accuracy onto the corresponding fUSI anatomical plane, keeping only the top 5% most significant voxels.

### Representational Similarity Analysis (RSA)

RSA was used to compare the relative similarity of each effector’s representation relative to all the others’. A representational dissimilarity matrix (RDM) was used to illustrate differences in representation across conditions. To generate RDMs, we computed the pairwise Crossnobis distances for each finger, averaged across sessions, as our dissimilarity metric. Crossnobis distance is a cross-validated, unbiased estimator of the squared Mahalanobis distance, a representation of the distance between two population-level response patterns^33^. Crossnobis distances were calculated for each BA subregion using the trial-averaged signal per voxel at peak activation for each effector and each run. The noise covariance matrix was estimated from the residuals of the GLM fit, which were temporally averaged in non-overlapping windows to account for temporal autocorrelation in the hemodynamic signal. Crossnobis distances were then computed using run-based split-half cross-validation. Finally, distances were normalized by the number of voxels per BA to account for Crossnobis distances scaling by number of voxels. Statistical significance of individual Crossnobis distances was assessed using permutation testing by randomly shuffling effector labels across trials and recomputing distances over 1000 permutations to generate a null distribution. Individual RDMs were calculated and visualized for each session in Figure S5.

## Acknowledgments

We thank our study participant, J, for their diligence and devotion to the study, which made this work possible. We thank Sunho Lee for their helpful discussions and insights. We thank Viktor Scherbatyuk for technical assistance.

## Funding

This research was supported by the National Institutes of Health (NIH) BRAIN Initiative (NIH R01NS123663 to R.A.A., M.G.S., and M.T.), the National Institute Of Neurological Disorders And Stroke (NINDS) of the NIH under the F30 (NINDS F30NS145610 to L.J.L.), the Merkin Institute for Translational Research (to R.A.A.), the Tianqiao and Chrissy Chen Brain-Machine Interface Center (to R.A.A. and M.G.S.), and the Boswell Foundation (to R.A.A.). The content is solely the responsibility of the authors and does not necessarily represent the official views of the NIH.

## Author contributions

L.J.L., T.C., M.G.S., and R.A.A. conceived the study. L.J.L. acquired the data. L.J.L. performed the data processing and analysis; B.H. assisted with data processing. K.P. provided administrative assistance and participant planning. C.L. conducted craniectomy and cranioplasty surgeries. L.J.L. drafted the manuscript with substantial contributions from T.C., B.H., M.G.S, and R.A.A., and all authors edited and approved the final version of the manuscript. C.L., M.G.S., and R.A.A. supervised the research.

## Competing Interests

M.G.S. is a cofounder and shareholder of Merge Labs. All other authors have declared no competing interest.

## Supplementary Figures

**Fig. S1:**
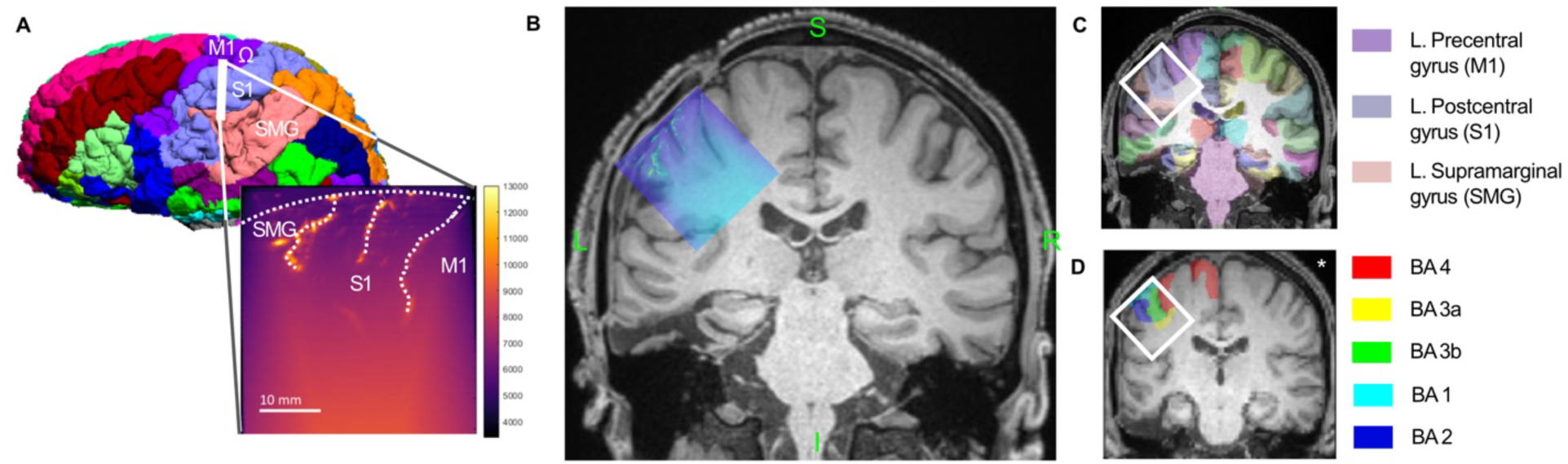
Manual alignment of fUSI plane to anatomical MRI. A) 3D rendering of the participant’s anatomical T1w MRI segmented into brain regions using BrainSuiteAtlas1 with the approximated fUSI plane location superimposed. The fUSI plane provided access to portions of SMG, S1, and M1 (white dotted lines represent cortical surface/sulci). B) fUSI plane aligned to a coronal section of participant’s MRI. Large vascular fiducials visualized in the fUSI plane were matched to the outline of sulci and gyri in the MRI. C) MRI coronal section overlayed with BrainSuiteAtlas1 anatomical labels. The white bounding box represents the approximate location of the fUSI plane relative to the labeled section. D) MRI coronal section overlayed with SPM25 Anatomy Toolbox cytoarchitectonic probability maps for BA 2, 1, 3b, 3a, and 4. The white bounding box represents the approximate location of the fUSI plane relative to the labeled section. (* = MRI was warped into MNI space to align with the probability maps).

**Fig. S2:**
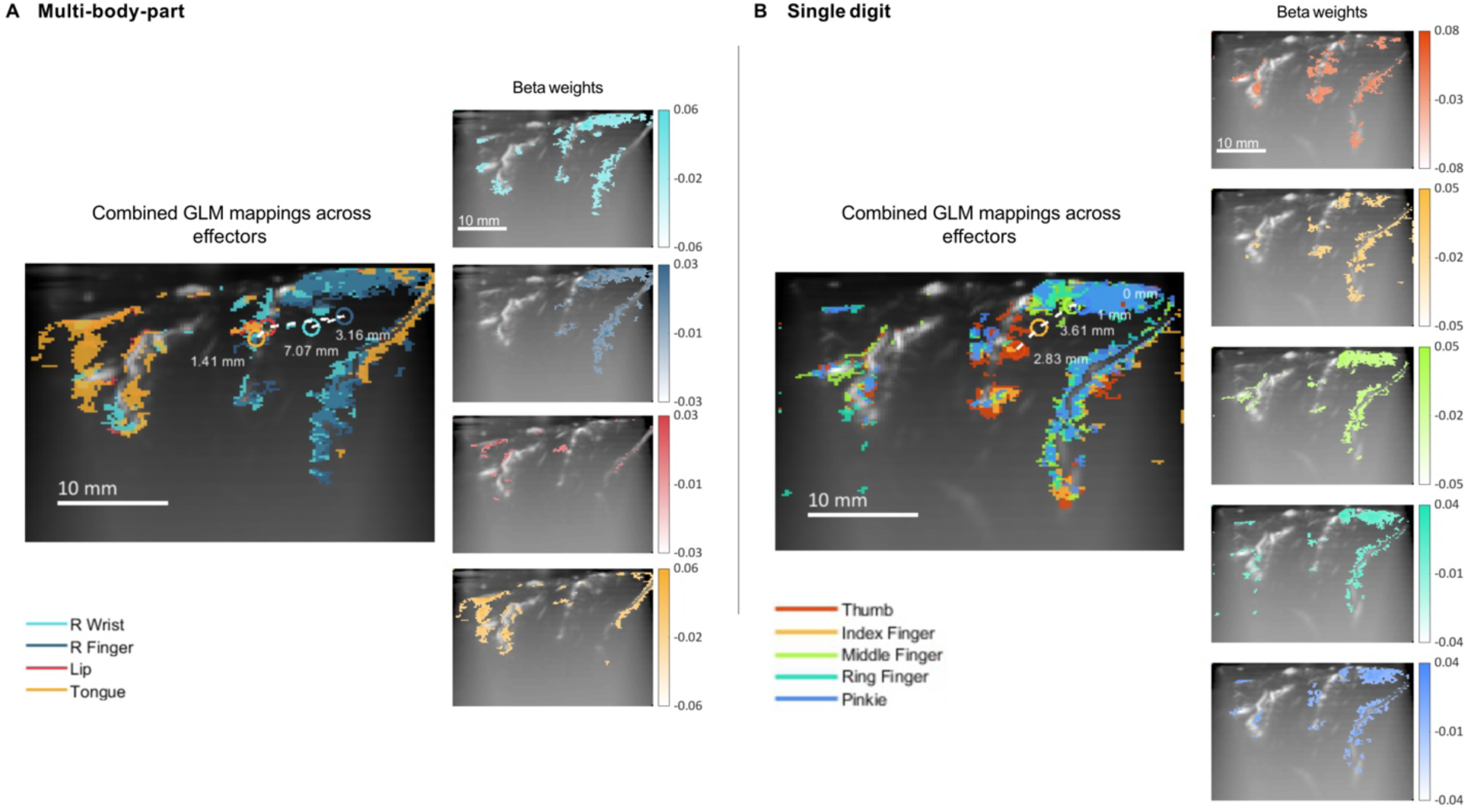
GLM maps of effectors relative to rest. A) Map of the combined statistically significant ROIs generated from GLM for multi-body-part effectors (compared to rest) and the individual ROIs per effector and their corresponding beta weights (session B1). The centroids of activation per ROI were calculated based on the regions of activation per condition within the S1 region and are indicated by a colored ring corresponding to each effector. The white dotted lines between centroids represent the connections between adjacent effectors based on the somatotopic model, with calculated Euclidian distances between centroids being displayed by their corresponding lines (p<1e-3 w FDR correction). B) Map of the combined statistically significant ROIs generated from GLMs for individual fingers (compared to rest) and the ROIs per finger and their corresponding beta weights (session F1). The centroids of activation per ROI were calculated based on the regions of activation per condition within the S1 region and are indicated by a colored ring corresponding to each effector. The white dotted lines between centroids represent the connections between adjacent effectors based on the somatotopic model, with calculated Euclidian distances between centroids being displayed by their corresponding lines (p<1e-3 w FDR correction).

**Fig. S3:**
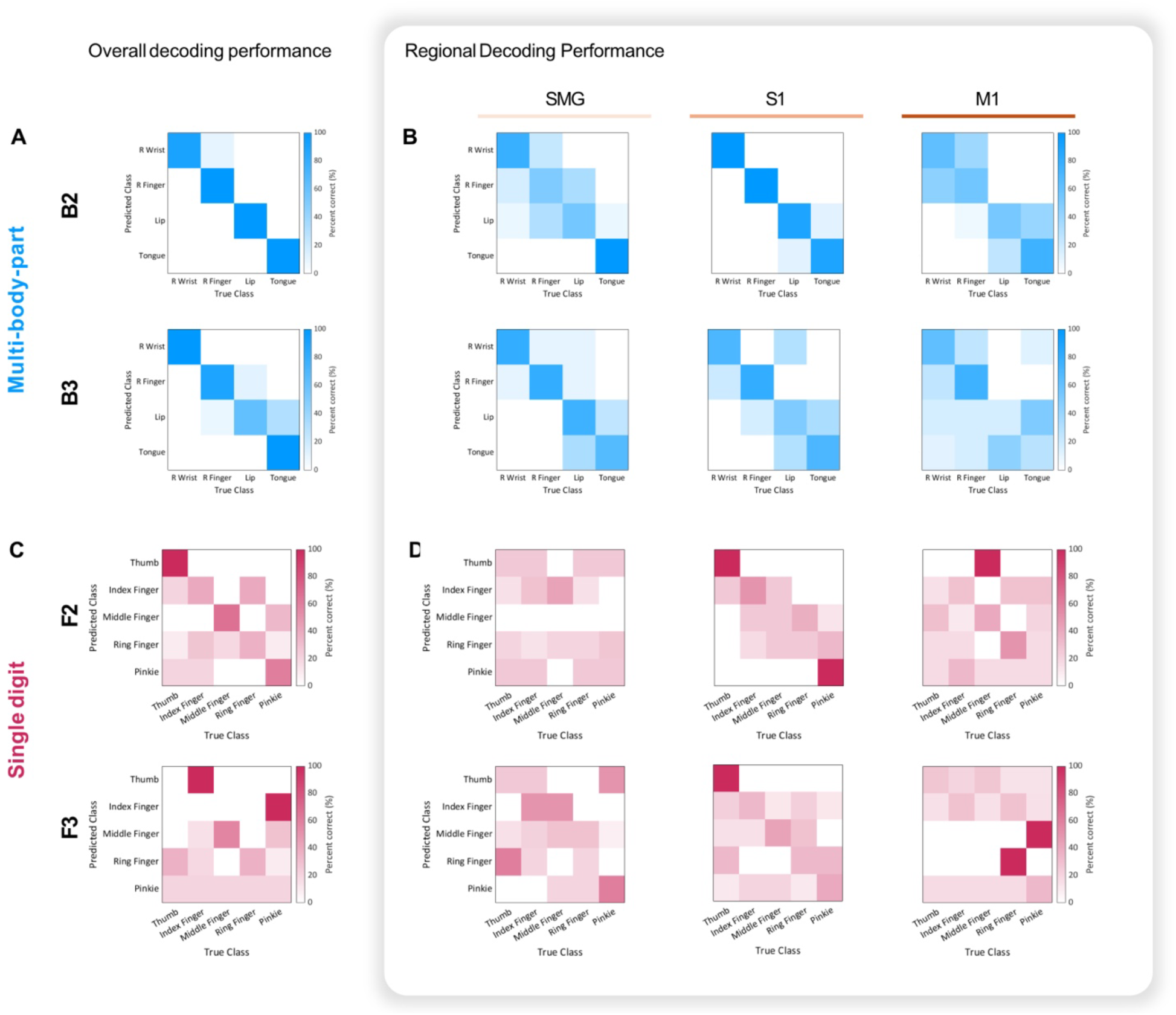
Additional session decoding results. A) Additional session decoding results from sessions B2 and B3 for multi-body-part movement. B) Additional session decoding results from sessions F2 and F3 for single digit movement.

**Fig. S4:**
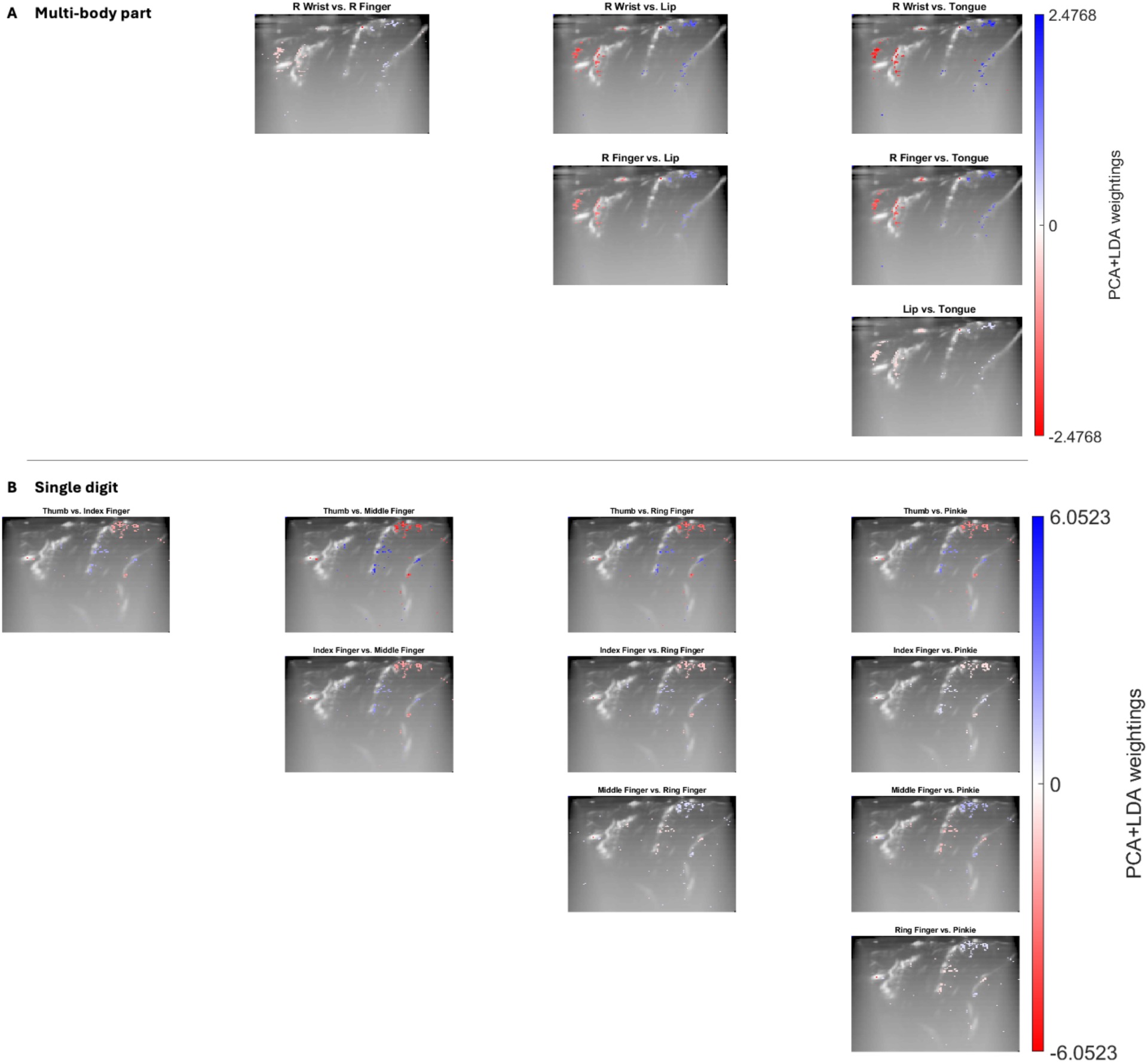
PCA-LDA weight maps. A) Multi-body-part PCA-LDA decoder weight maps from session B1. LDA weights of PCA features were taken from a PCA-LDA decoder trained on the entire fUSI image at the timepoint with the best decoding accuracy. The PCA features were then projected back into voxel space and overlayed on anatomical fUSI images. Highlighted regions represent the top 1% highest weighted voxels for distinguishing the labeled pair of effectors. B) Single digit PCA-LDA decoder weight maps from session F1. Highlighted regions represent the top 1% highest weighted voxels for distinguishing the labeled pair of effectors.

**Fig. S5:**
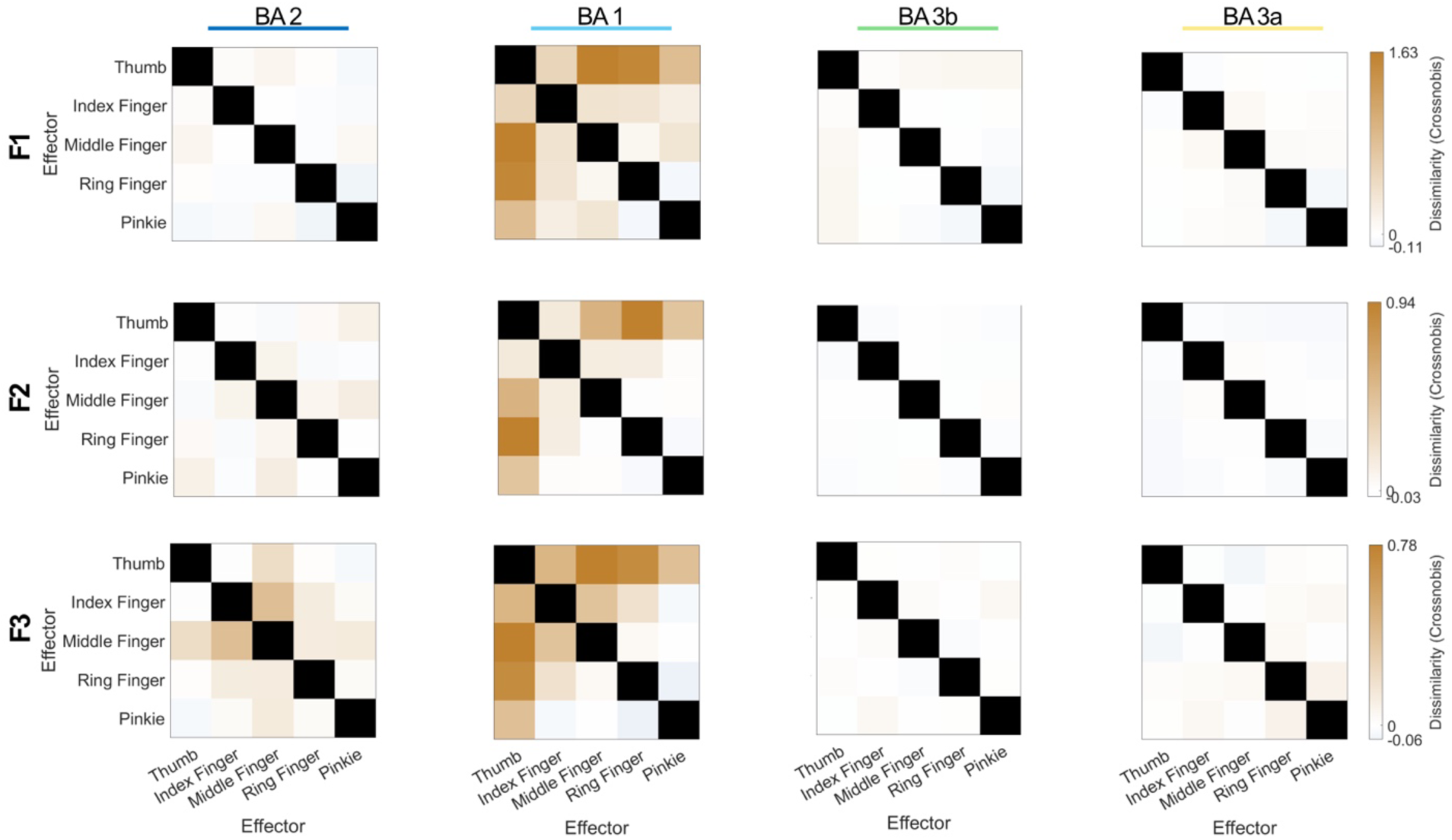
RDMs across cytoarchitectonic regions across individual sessions. RDMs generated for each BA per session (session F1, F2, F3) to compare representational differences across cytoarchitectonic regions. RDMs were generated by calculating the Crossnobis distances from trial-averaged signal per voxel at peak activation for each effector and each run.

**Fig. S6:**
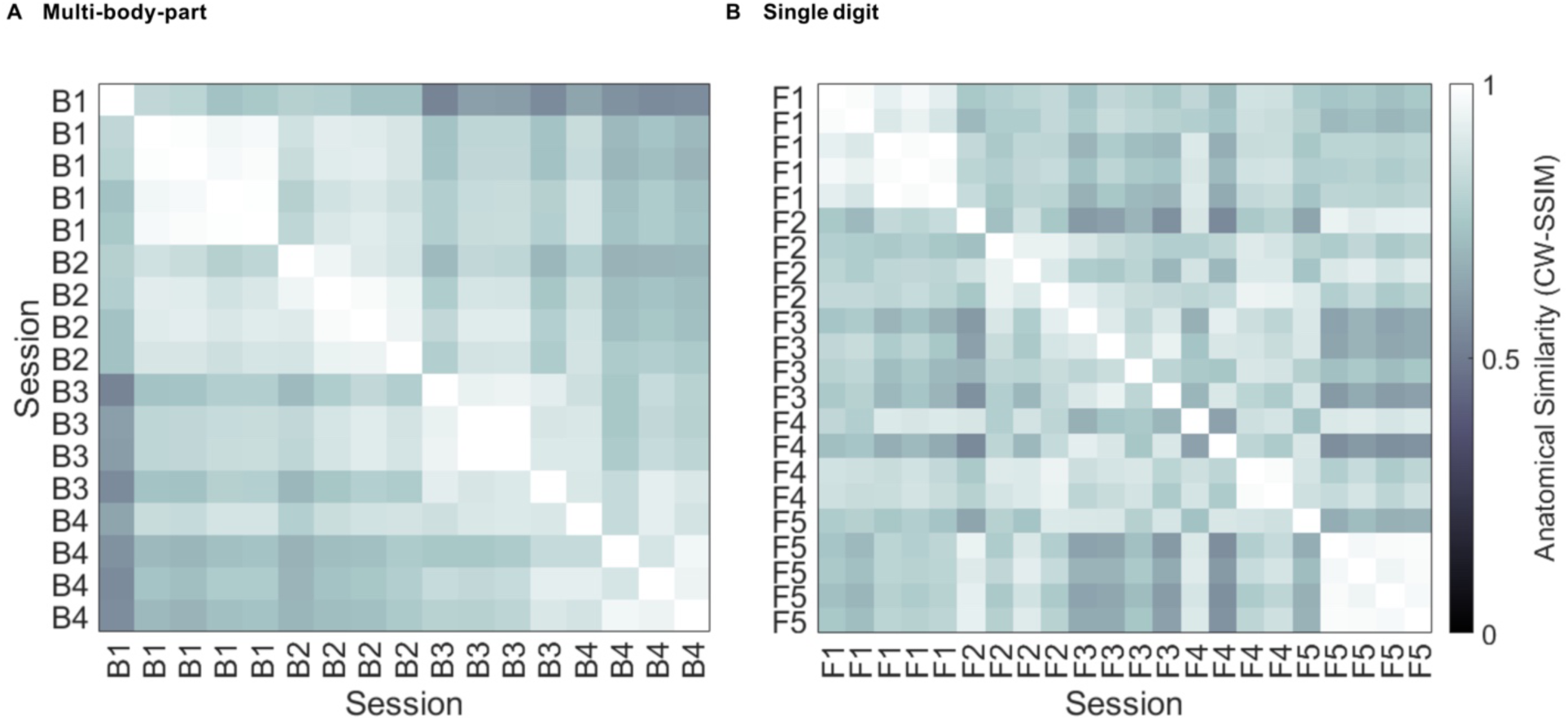
Pairwise CW-SSIM values across all experiment runs. The pairwise CW-SSIM was calculated for all run pairs to determine the similarity of imaging planes across runs. Runs where CW-SSIM < 0.6 relative to runs from sessions B1 and F1 were deemed as different planes and excluded. A) Multi-body-part movement. B) Single digit movement.

